# Phylo-rs: an extensible phylogenetic analysis library in Rust

**DOI:** 10.1101/2025.03.10.642340

**Authors:** Sriram Vijendran, Tavis K. Anderson, Alexey Markin, Oliver Eulenstein

## Abstract

We introduce Phylo-rs: a fast, extensible, general-purpose library for phylogenetic analysis and inference written in the Rust programming language. Phylo-rs leverages a combination of speed, memory-safety, and native WebAssembly support offered by Rust to provide a robust set of memory-efficient data structures and elementary phylogenetic algorithms. Phylo-rs is focused on efficient and convenient deployment of software aimed at large-scale phylogenetic analysis and inference. Phylo-rs is available under an open-source license on GitHub at https://github.com/sriram98v/phylo-rs, with documentation available at https://docs.rs/phylo/latest/phylo/.

## Introduction

Phylogenetic trees, or *phylogenies*, are fundamental to evolutionary biology as they represent hypotheses about the relationships between different taxonomic groups, benefiting diverse disciplines from agronomy [1] and conservation biology [2–4] to medical sciences [5] and epidemiology [6]. Recent advances in next-generation and long-read sequencing technologies [7, 8] have improved access to large-scale genomic data and phylogenies. The scale of these data and phylogenetic trees necessitates efficient and effective computational libraries that implement specialized algorithms to analyze phylogenies and uncover hidden statistics and relationships between taxonomic groups [9–11].

Current phylogenetic libraries have, at times, struggled to keep pace with the demands of large-scale phylogenetic analysis. Existing libraries often make trade-offs between runtime efficiency and developmental ease based on the chosen language. Software implemented in libraries like Dendropy [12], TreeSwift [13], phytools [14] and ape [15] offer simple and intuitive syntax at the cost of the efficiency, low-level control, and functionality necessary for large-scale phylogenetic analysis. In contrast, implementations in libraries like Bio++ [16] and Gotree [17] offer memory and runtime efficiency but lack the memory-safety and security features of modern programming languages [18, 19].

Rust is a modern programming language that leverages speed and memory-safety with high-level syntactical features. Rust is compiled with LLVM [20], providing optimal speed with a low memory footprint. Additionally, Rust supports automatic type inference at compile time, reducing the verbosity of written code. The key feature of Rust is the concept of ownership and borrowing of variables, which enables Rust to infer the lifetime of data stored in memory automatically. This eliminates the overhead of online memory management and completely eradicates common memory errors such as segmentation faults. Concomitantly, ownership enforces thread-safety, preventing race conditions in multi-threaded code. These features make Rust an attractive alternative for applications in Bioinformatics.

We introduce Phylo-rs, a versatile phylogenetic library that provides an extensible foundation of data structures and algorithms for phylogenetic analysis and inference implemented in Rust [21]. Phylo-rs utilizes Rust’s modern programming language features, delivering high-performance software while ensuring memory-safety and maintainable code. Additionally, Phylo-rs provides native WebAssembly (WASM) support, offering a highly portable and compact compilation target for software [22]. This enables access to software written using Phylo-rs on web browsers, eliminating system compatibility issues and narrowing the gap between cutting-edge research and practical application [23]. To our knowledge, Phylo-rs is the first comprehensive phylogenetic analysis library written in Rust.

## Design and Implementation

At a high level, phylogenies in Phylo-rs are implemented as Rust ‘traits’ that describe their behavior and functionality while making no assumptions on how they are represented in memory. These traits allow using any data structure, also called *structs*, to represent phylogenies. Structs require the implementation of only a few basic methods to gain access to several iterators, operators, and functions. This includes tree traversals, simulations, distance metrics, edit operations, and file I/O. These traits can be inherited by other user-defined traits, enabling seamless extensions to existing methods and convenient implementation of new algorithms, as shown in Figure 1.

**Figure 1.**
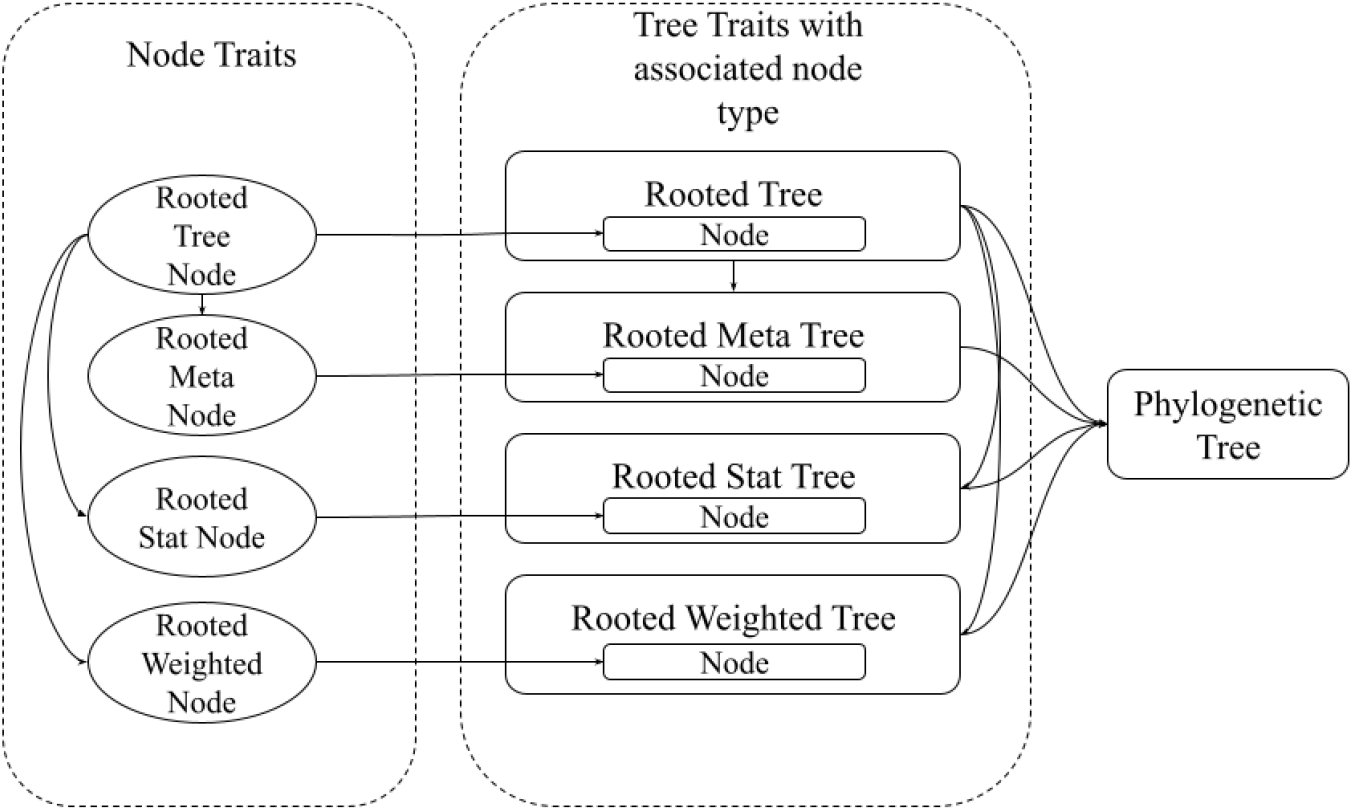
A trait dependency graph showing how behavior is shared between objects that build up to a phylogenetic tree. Meta tree nodes, stat tree nodes, and weighted tree nodes extend the behavior of a rooted tree node to manipulate the meta-annotation, stat-annotation, and weight-annotation of a node, respectively. Similarly, a rooted meta tree and a rooted stat tree extend the behaviors of a rooted tree, and finally, a phylogenetic tree extends the behavior of all the defined trees

Phylo-rs eliminates redundant memory usage by yielding references instead of deep copies. Phylo-rs enforces memory-safety at compilation, which secures software from memory vulnerabilities. Memory-safety is ensured in Phylo-rs by assigning object lifetimes; tree components are retained in memory for as long as the tree itself, eliminating memory-related errors or vulnerabilities.

Classical analyses of phylogenies require the pairwise comparison of trees using established metrics such as the Robinson-Foulds metric [24], cophenetic distances [25, 26], and cluster affinity distance [27]. Phylo-rs offers functions that implement the most efficient known algorithm [28] to compute these distances.

Many phylogenetic inference algorithms employ tree edit operations [29–32] in algorithms aimed at inferring the optimal phylogenetic history of a set of taxa. In line with that, Phylo-rs provides traits to perform tree edit operations such as Subtree Pruning and Regrafting [33], Tree Bisection and Reconnection [34], and Nearest Neighbor Interchange [35].

Phylo-rs supports the widely used Newick [36] encoding for phylogenies, including constructing and translating trees from live streams of ASCII data over web-based and multi-threaded ports. Phylo-rs implements a Newick trait that can be extended to cloud-based applications. The Newick trait can also be extended to support numerous file formats, such as the Nexus format, without making any metadata structure specifications.

Phylo-rs is furnished with an intuitive tree-like struct that implements all the traits of phylogenies, which is fully detailed in the official Phylo-rs documentation. Phylo-rs documents the trade-offs for every method, providing links to alternative methods that achieve the same results differently, where possible. Traits are automatically tested using the standard tree struct via continuous integration and are benchmarked at every stable release.

Phylo-rs is equipped with additional features to enable researchers to implement algorithms for large-scale analysis seamlessly. Each feature can be enabled or disabled at compilation time, depending on the infrastructure of the target hardware.

### Multi-threading

Phylo-rs delivers multi-thread support by parallelizing its iterators while guaranteeing data-race freedom. Analyses that require independent computations for each vertex of a phylogeny can be executed simultaneously. Data parallelism can be highly beneficial in large-scale studies where phylogenies with tens of thousands of taxa can be analyzed efficiently by sharing the computational workload between numerous CPUs.

### Single Instruction, Multiple Data

Phylo-rs permits parallelization of bit-level operations on single-CPU environments through the use of Single Instruction, Multiple Data (SIMD). SIMD has been frequently used to improve application performance in a variety of fields [37–39], with cases achieving a 9x speedup [40]. Phylo-rs utilizes SIMD when inferring and enumerating bipartitions of the taxa induced by a phylogeny. Phylo-rs computes the overlap between two clusters through parallelized bit-level operations on the same core by representing clusters as bit-strings.

### WASM

Phylo-rs achieves platform interoperability, ease of use, and effortless distribution by supporting WASM as a compilation target. WASM is a compact binary instruction format for stack-based virtual machines [22] and can be called from JavaScript via Node.js or as a command line interface application. With WASM support, Phylo-rs imparts:

Safety: Users are protected by software sandboxed virtual environments, protecting them from any damage from running malicious code.

Speed: Low-level code generated by compilers is optimized ahead of time, allowing the code to fully utilize machine hardware; WASM supplies users with efficient tools that overcome the inefficient runtimes traditionally seen with sandboxed applications.

Portability: Low-level code compiled to WASM as a single architecture targeted for the Web can run across various browsers, operating systems, and hardware types.

As such, WASM is an excellent alternative to standard Graphical User Interface applications and provides a robust platform for disseminating bioinformatic tools and applications [41, 42]. User interfaces can be standardized using any modern web browser, reducing the redundant graphical overhead of installed applications. Analytic tools written with Phylo-rs can be shared as web apps with built-in graphical interfaces and intuitive visualizations using modern graphical libraries [43].

## Results

We present a comparative analysis highlighting the performance of Phylo-rs relative to popular libraries, namely, Dendropy [12], GoTree [17], TreeSwift [13], and ape [15]. Following this, we demonstrate its utility with two examples of computationally demanding phylogenetic analyses that can be solved using Phylo-rs. All results and corresponding visualizations presented in this section can be reproduced on a typical desktop PC by following the instructions in the official GitHub repository at https://github.com/sriram98v/phylo-rs.

### Comparative Analysis

We compare Phylo-rs with other popular phylogenetic libraries using a benchmark study that contrasts the mean runtime of six foundational algorithms commonly employed in phylogenetic analyses [29, 30]: (i) computing the Robinson-Foulds metric, (ii) retrieving the Least Common Ancestor (LCA), (iii) tree traversals in pre- and post-order for vertices and edges, (iv) subtree extraction and contraction, (v) simulating random trees using the Yule evolutionary model, and (vi) applying the Nearest Neighbor Interchange (NNI) operation.

We conducted 1000 iterations for each implementation with a precision of ±12 ns on a randomly simulated phylogenetic tree comprising 4000 taxa, excluding libraries that did not provide an implementation. When evaluating the computation of the Robinson-Foulds metric, we measured 1000 iterations on a pair of randomly simulated trees. We assessed tree simulation by measuring 1000 iterations of generating a tree comprising 4000 taxa. All benchmarks were conducted with *Cargo bench* for Phylo-rs, Python *timeit* for Dendropy [12] and TreeSwift [13], and the Linux *time* utility for ape [15] and Gotree [17], using identical trees for each implementation of the same algorithm. Each benchmark was performed on an Intel(R) Core(TM) i7-10700K 3.80GHz CPU running Arch Linux v6.6.28-2-lts and was executed on a single thread. Benchmarking with *timeit* and *time* entails an overhead of approximately 2 ms for reading the trees from files, which was excluded from the recorded runtimes.

#### Benchmark Study

Table 1 summarizes the mean runtime of computing the Robinson-Foulds metric, tree contraction, tree traversal, Yule tree simulation, LCA retrieval, and NNI. Notably, Phylo-rs achieves a significant speedup compared to the libraries written in Python and R while showing comparable performance with Gotree [17], written in Go-lang; Phylo-rs achieves a 10x speedup compared to Dendropy [12] and ape [15] on all compared implementations and a nearly 2x speedup in computation compared to GoTree. These operations are fundamental components of many popular algorithms used in practice, including maximum likelihood estimation [29, 30] and Bayesian inference [44]. The improved runtimes indicate that Phylo-rs can significantly reduce the time required to perform large-scale phylogenetic analyses, making it a more efficient choice for researchers and practitioners. Furthermore, the comparable performance of Phylo-rs with GoTree in tree traversal and LCA retrieval suggests that Phylo-rs is a viable alternative for existing phylogenetic analysis workflows.

**Table 1.** List of algorithms commonly used in computational phylogenetics.

| Algorithms | Phylo-rs | Dendropy | GoTree | TreeSwift | ape |
| --- | --- | --- | --- | --- | --- |
| Subtree Prune-Regraft<sup>a</sup> | 2.01 ns | - | - | - | - |
| Tree Bisection-Reconnection<sup>a</sup> | 2.01 ns | - | - | - | - |
| Nearest Neighbor Interchange<sup>a</sup> | 2.01 ns | - | 4.02 ns | - | - |
| Robinson-Foulds metric | 16.53 ms | 160.80 ms | 25.02 ms | - | 250.14 ms |
| cluster matching distance | 88.52 ms | - | - | - | - |
| LCA | 5.4 ms | 78.10 ms | - | 11.54 ms | 327.81 ms |
| Tree Contraction | 2.45 ms | 71.27 ms | 3.87 ms | 11.56 ms | - |
| Vertex Traversal<sup>b</sup> | 0.34 ms | 61.29 ms | 0.56 ms | 10.96 ms | 50.21 ms |
| Edge Traversal<sup>b</sup> | 0.34 ms | 61.29 ms | 0.56 ms | 10.96 ms | 50.21 ms |
| Yule Tree Simulation | 32.78 ms | -* | 65.17 ms | - | - |

<sup>a</sup> and <sup>b</sup> have identical runtimes as they all use the same underlying implementation in Phylo-rs.
\* This result has been omitted as Phylo-rs simulates only topology while Dendropy simulates topology and branch lengths, which adds a significant computational overhead.

Additionally, Table 1 indicates that there are more methods natively implemented in Phylo-rs than those compared in the previous section. These additional operations are essential for many applications in phylogenetic analysis, such as tree reconciliation and phylogenetic inference. Phylo-rs can be easily integrated into various workflows and pipelines by providing a broader range of fundamental operations. This makes it appealing for researchers and practitioners working on diverse phylogenetic tasks.

### Quantifying Phylogenetic Diversity for Influenza A Virus Control

We quantified the phylogenetic diversity (PD) [45] of the H1 subtype influenza A virus (IAV) in swine collected between the years 2015 and 2022. The H1 subtype of swine IAV in the United States has at least 11 genetically distinct clades of viruses [46]. Controlling IAV transmission relies upon vaccination and designing optimal vaccination strategies requires a detailed analysis of the genetic diversity of the circulating viruses [47, 48].

To quantify diversity dynamics, we downloaded all 8241 publicly available IAV hemagglutinin (HA) sequences from the USDA influenza A virus in the swine surveillance system collected between 2015–2022. All sequences were classified into one of the named swine IAV clades using octoFLU v.1.0.0 [46, 49]. We aligned the nucleotide sequences with mafft v.7.525 [50] and inferred a maximum likelihood tree using IQ-Tree v2.2.6 [30] under the generalized time-reversible (GTR) substitution model [51] with empirical base frequencies and five free-rate categories [52]. We computed PD for each named clade detected within each year using Phylo-rs and visualized the resulting dynamics in Figure 2. These data indicate that the 1B.2.1 and 1A.1.1.3 clades demonstrated a steady increase in PD across the years, whereas other clades, e.g., 1B.2.2.1 and 1A.3.3.2, fluctuated. The steady increase in PD in the 1B.2.1 and 1A.1.1.3 clades represents a significant challenge for control strategies, i.e., vaccines to reflect circulating genetic and antigenic diversity may not work adequately as a strain selected as a vaccine antigen in 2016 may not reflect the diversity in the clade in 2018 [48]. In addition, this analysis identified clades with low PD, which may be susceptible to removal through the use of targeted vaccines that are focused on the genetic diversity observed within these clades. A benefit of using PD to track diversity is that clades may be driven to extinction with a reduction in total genetic diversity and the subsequent minimization of reassortment and antigenic drift [53].

**Figure 2.**
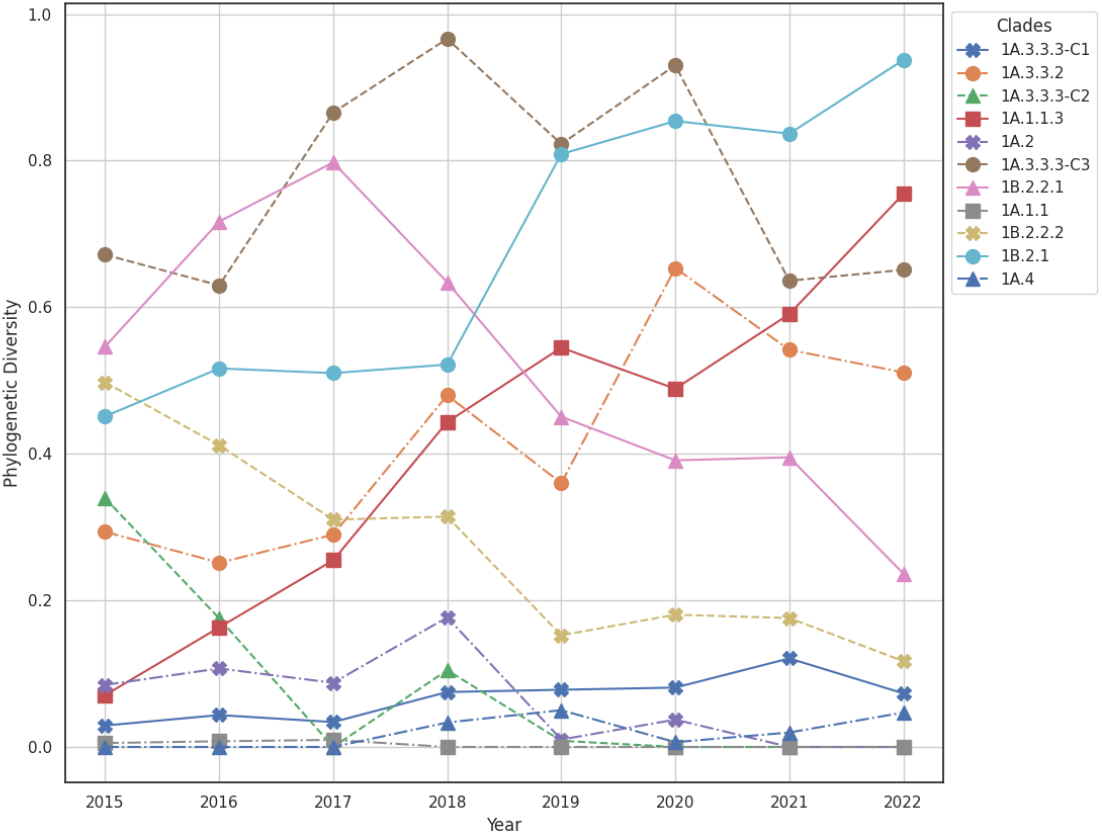
Visualization of variation in phylogenetic diversity of the H1 subtype influenza A virus (IAV) collected between the years 2015 and 2024. The phylogenetic clades 1B.2.1 and 1A.1.1.3 demonstrated an almost linear increase in phylogenetic diversity across the years indicating evolution of the pathogen with increases in genetic diversity that may reduce the efficacy of vaccine control strategies. The phylogenetic clades 1B.2.2.2 and 1A.4 demonstrated a decline in phylogenetic diversity, suggesting that vaccine control measures may be designed with a single antigenic component to effectively prevent infection and transmission. The phylogenetic diversity of a tree at each year was computed as the Faith Index [45] implemented in Phylo-rs.

### Visualizing Phylogenetic Tree Space

Phylogenetic tree spaces are often complex with many local optima, which confounds the phylogenetic inference [29, 30, 44]. A standard approach to searching the tree space for an optimal phylogeny is to sample the tree space using multiple Markov chain Monte Carlo (MCMC) Bayesian analyses [29, 30, 44], resulting in several samples of the tree space. The samples produced by each analysis can then be visualized by computing all pairwise distances between the sampled trees and embedding them into a 2- or dimensional Euclidean space [54]. A single MCMC analysis can produce upwards of 10000 trees, making the computation of pairwise distances infeasible in large-scale studies involving hundreds of taxa. Phylo-rs makes the computation of all pairwise distances feasible even on large datasets with thousands of taxa and tens of thousands of sampled trees due to its innate speed and in-built multi-threading.

We tested this approach on a MCMC analysis that was conducted to assess the emergence and spread of highly pathogenic avian influenza (HPAI) H5N1 viruses in dairy cattle in the US from [55]. Ten independent MCMC runs were conducted with BEAST v1.10.4 on a set of 587 influenza A virus hemagglutinin H5N1 clade 2.3.4.4b sequences sampled from dairy cattle, poultry, peridomestic mammals, and wild birds. Each run consisted of a single Markov chain lasting 50 million generations, sampled every 5000 steps. This resulted in 10001 sampled trees in each run and 100010 trees in total. We computed all pairwise Robinson-Foulds metrics between the sampled trees using Phylo-rs on a workstation with an Intel(R) Xenon(R) w7-2475X 4.8GHz CPU running Ubuntu 20.04.3 LTS. The computation was conducted with 40 threads, taking 32 hours to calculate the distance of approximately 5 billion tree pairs.

To simplify visualization, we omitted four runs that did not converge [55] and removed the first 20% of trees as the burn-in from the remaining 6 runs. We then embedded the distances between the remaining 48,000 trees into a 2-dimensional space using UMAP (Figure 3). Each independent MCMC run formed a continuous line in the resulting embedding. All runs except for run 10 appear to have traversed a similar subspace of trees while run 10 clusters separately from the other runs.

**Figure 3.**
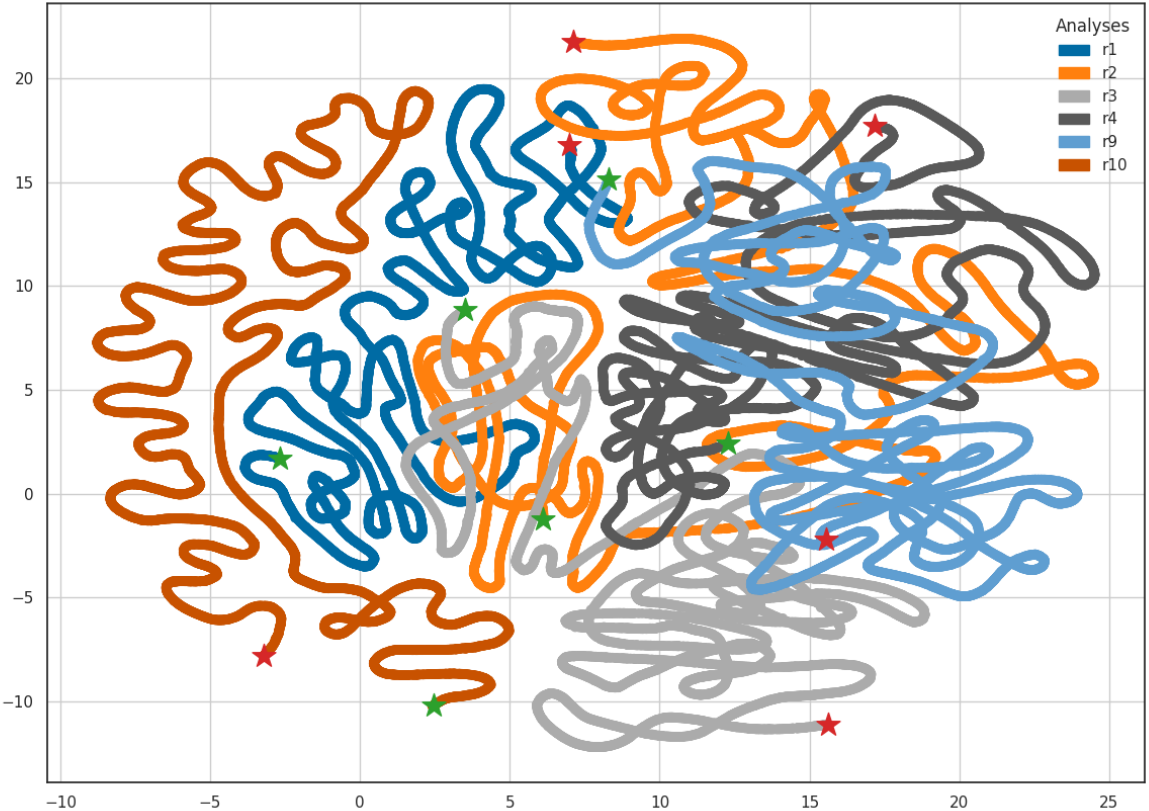
UMAP embedding of the phylogenetic tree space explored by 6 independent MCMC runs. All runs were conducted under the same conditions. Each color represents the trees from a single run, where the green star indicates the starting tree and the red star indicates the final tree. The distances between the trees were computed using the Robinson-Foulds metric as implemented in Phylo-rs.

## Availability and Future Work

Phylo-rs is a general-purpose phylogenetic analysis library written in Rust. By leveraging the Rust programming language’s memory-safety features and speed, Phylo-rs offers a variety of advanced phylogenetic algorithms and functionality. Phylo-rs fosters the dissemination of complex software for phylogenetic analysis, bridging the gap between theoretical advancement and practical implementation. Phylo-rs is available under an open-source license on GitHub at

https://github.com/sriram98v/phylo-rs, with documentation at

https://docs.rs/phylo/latest/phylo/.

Support for PhyloXML and PhyloJSON file formats will be included in the future. Further, tree simulations under the Birth-Death and Coalescent evolutionary models will in added in the near future. Phylo-rs will extend bindings to other languages, such as R and Python, and implement tree traits on highly memory-efficient structures provided by libraries such as ts-kit [56].

## Supporting information

This project was funded in part by the United States Department of Agriculture (USDA), Agricultural Research Service (ARS project numbers 5030-32000-231-000-D, 5030-32000-231-111-I, 3022-32000-018-017-S, 5030-32000-231-095-S, and 5030-32000-231-103-A) and with federal funds from the National Institute of Allergy and Infectious Diseases, National Institutes of Health, Department of Health and Human Services (Contract No. 75N93021C00015). The funding sources had no role in study design, data collection, and interpretation, or the decision to submit the work for publication. Mention of trade names or commercial products in this article is solely to provide specific information and does not imply recommendation or endorsement by the USDA. USDA is an equal opportunity provider and employer.

## Acknowledgments

We are grateful for the comments on the manuscript provided by Dr. Pawewl Górecki, Dr. Geng Ding, and Paige Falor and code reviews by Sanket Wagle.

